# Directed connectomes across species reveal conserved and divergent pathways of neural signaling

**DOI:** 10.1101/2025.09.07.674762

**Authors:** Siva Venkadesh, Wen-Jieh Linn, Yuhe Tian, G Allan Johnson, Fang-Cheng Yeh

## Abstract

Understanding how brain networks support cognition requires knowing not only which regions are connected but the direction in which neuronal signals propagate. However, directed connectivity cannot be measured non-invasively in primates or humans, limiting systems-level inference. Here we integrate projection polarity from ∼1,200 mouse viral tracing experiments with species-specific diffusion MRI tractography to construct directed connectomes across mouse, marmoset, macaque, and human. Using a common cross-species atlas, we developed a path efficiency metric that balances projection strength against axonal length and applied shortest-path algorithms to quantify directional influence. This framework revealed conserved and divergent organization: the entorhinal–hippocampal projection was the most efficient in all species; humans showed strengthened anterior insula–superior temporal pathways; macaques exhibited peak inferior temporal outflows; and marmosets maintained disproportionately strong olfactory influence. These results provide a scalable and ethically feasible approach for modeling directed neuronal signaling across species.

## 1. Introduction

Understanding how brain networks are conserved across species, and how they diverge to support lineage-specific functions, is a central goal of translational neuroscience. Neural systems evolve under selection pressures and limited resources, so expansion in one domain can entail reorganization elsewhere, producing trade-offs in specialization^1,2^. Complex brain functions emerge from the unidirectional flow of neuronal signals through large-scale pathways, making the reconstruction of *directed* networks essential for understanding the organizational principles of brain connectivity. Applications of graph theory and dynamical systems models of network function depend critically on resolving the directionality of connections^3^, which reflect the anatomical polarity of neuronal projections between regions: signals originate at the neuronal soma, travel along the axon, and terminate at synapses onto downstream targets. Viral tract-tracing provides gold-standard evidence for this projection polarity, but its invasive nature restricts application to small-brained species such as the mouse^4^. By contrast, diffusion MRI (dMRI) enables whole-brain mapping in primates and humans but yields bidirectional estimates that obscure projection polarity^3,5,6^. This methodological disconnect has limited our ability to test how conserved communication pathways coexist with lineage-specific specializations in association, sensory, and limbic systems.

Building on a previously developed common hierarchical atlas^7^, we combined the complementary strengths of viral tract-tracing and diffusion MRI. Viral tracing in the mouse brain, based on ∼1,200 anterograde injection experiments^4^, provides high-resolution, empirically validated directional polarity of axonal projections, while dMRI offers whole-brain coverage in humans and primates without directional specificity. By combining tracer-derived directionality with species-specific dMRI tractography, we constructed directed structural connectomes for each species. Brain circuitry, the pathways supporting the forward signal flow, is represented as a directed connectome with inter-regional connections modeled as asymmetric edges. We then quantified inter-regional influence using path efficiency, based on a novel multi-objective cost that favors shorter, stronger projections and reflects the physical and metabolic investment per unit influence. Directed shortest-path algorithms^8^ were applied to these networks, with results normalized for brain size and network scale to enable cross-species comparison.

Within a Darwinian framework, statistical differences across species reflect evolutionary pressures that have shaped network organization across lineages^9^. Our analysis revealed both conserved pathways and lineage-specific reallocations in cortical communication. Across all species, the entorhinal–hippocampal projection ranked as the most efficient, underscoring the evolutionary preservation of memory-related circuitry. In humans, directed paths involving the anterior insula and superior temporal cortex showed pronounced efficiency gains, highlighting a strengthened temporal–insula–frontal circuit central to multimodal integration and language-related functions. In macaques, inferior temporal outflows peaked, consistent with ventral visual stream specialization, while in marmosets, olfactory influence remained prominent relative to other primates. Together, these findings highlight conserved memory circuits and divergent association and sensory pathways, enabled by directionally informed connectomics in a common atlas scaffold.

## 2. Results

We leveraged a recently developed common cross-species atlas^7^ as the anatomical scaffold to integrate mouse viral tracing with diffusion MRI tractography and to generate cross-species brain circuitry (**Fig. 1**). Viral tracer experiments in the mouse (∼1,200 anterograde injections)^4^ provide empirically validated projection polarity, while dMRI supplies whole-brain trajectories and axonal lengths in mouse, marmoset, rhesus macaque, and human. By combining these complementary modalities, we constructed directed connectivity matrices in which inter-regional edges are asymmetric and weighted by an effective connection cost, a multi-objective metric that penalizes long, weak projections and favors short, strong ones. Directed shortest-path algorithms applied to these matrices identify the most efficient influence channels through which electrical activity can propagate.

**Figure 1.**
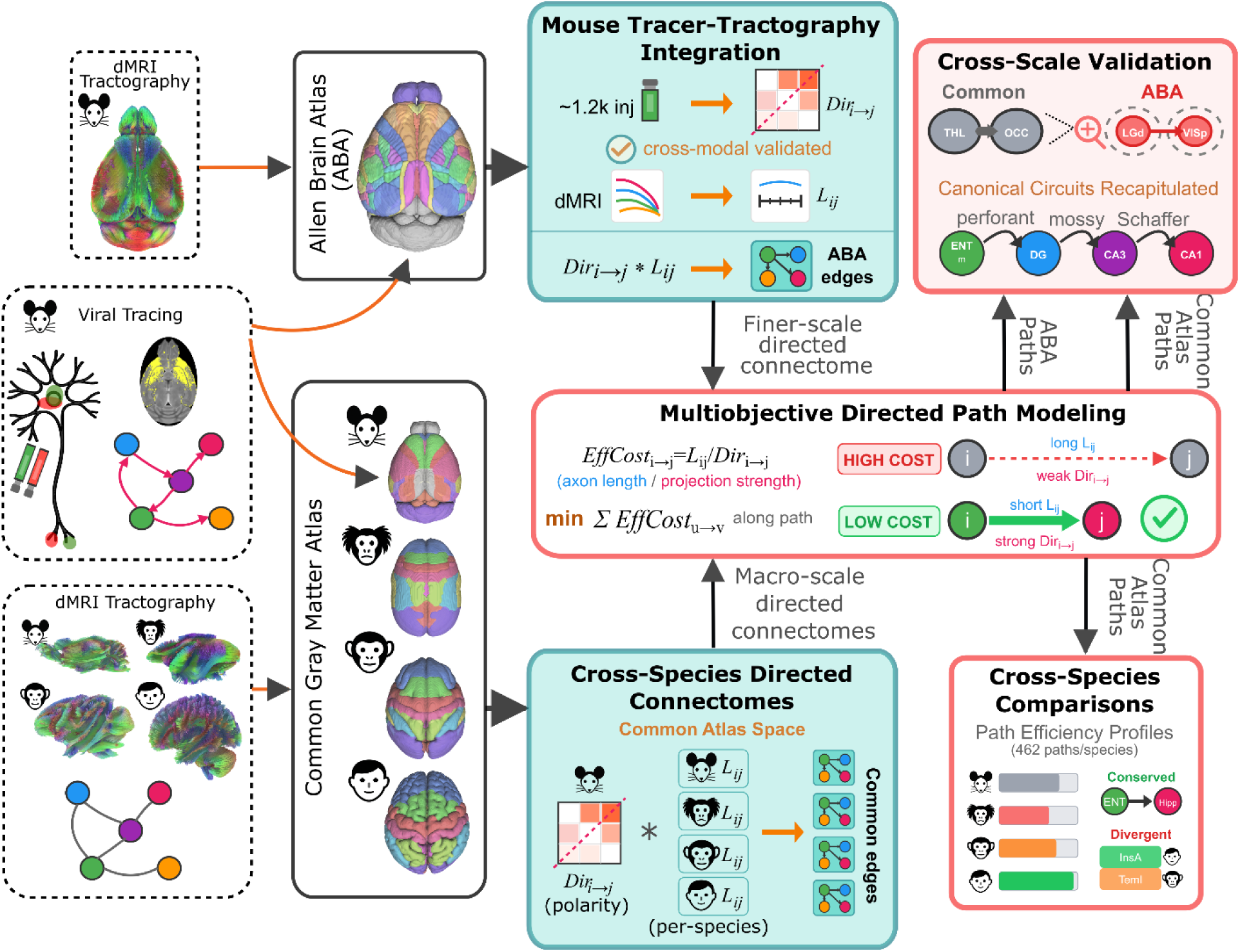
Framework for constructing cross-species directed connectomes. Mouse viral tracer data (∼1,200 anterograde injections) provide empirically validated projection polarity, which is mapped onto two anatomical scaffolds, the Allen Brain Atlas (ABA) for finer-scale analysis and a common cross-species atlas for macro-scale comparison, and combined with species-specific diffusion MRI tractography to generate directed connectomes in mouse, marmoset, rhesus macaque, and human. Effective connection cost, a multi-objective metric favoring shorter and stronger projections, is minimized along directed paths to identify efficient influence channels. Cross-scale validation tests correspondence between macro-level and finer-scale pathways. Cross-species comparisons quantify path efficiency profiles to identify conserved and divergent organization across species.

This framework enables analyses at multiple scales. At the cross-species level, the common atlas provides a consistent scaffold for identifying conserved versus lineage-specific reallocations of efficiency. At the fine scale, the same procedures can be applied within the Allen Brain Atlas (ABA), which serves two purposes: to recapitulate canonical mouse circuits and to validate that macro-level pathways identified in the common atlas are consistent with finer-scale organization. Together, these elements establish a scalable approach for directed connectomics that links tracer-derived polarity with tractography-derived wiring constraints, providing the foundation for the cross-species and cross-scale analyses that follow.

### 2.1. Cross-modal validation and construction of directed connectomes

We first validated the integration of viral tracing and diffusion MRI tractography in the mouse before extending the framework to construct directed connectomes across species. In the mouse, individual viral tracer injections covered on average ∼50% of an Allen Brain Atlas (ABA) ROI (**Fig. 2a**), underscoring the value of tractography for achieving near-complete sampling of white-matter projections. A representative tracer experiment is shown in **Fig. 2b**. Tractography seeded from injection sites reproduced more than 90% of seed-only tracts when constrained with tracer-defined projection endpoints, whereas random endpoints yielded far fewer matches (Wilcoxon signed-rank test, one-tailed, greater: W = 680,071, p < 0.0001, r = 0.80; **Fig. 2c**), confirming strong cross-modal correspondence consistent with a prior report^10^.

**Figure 2.**
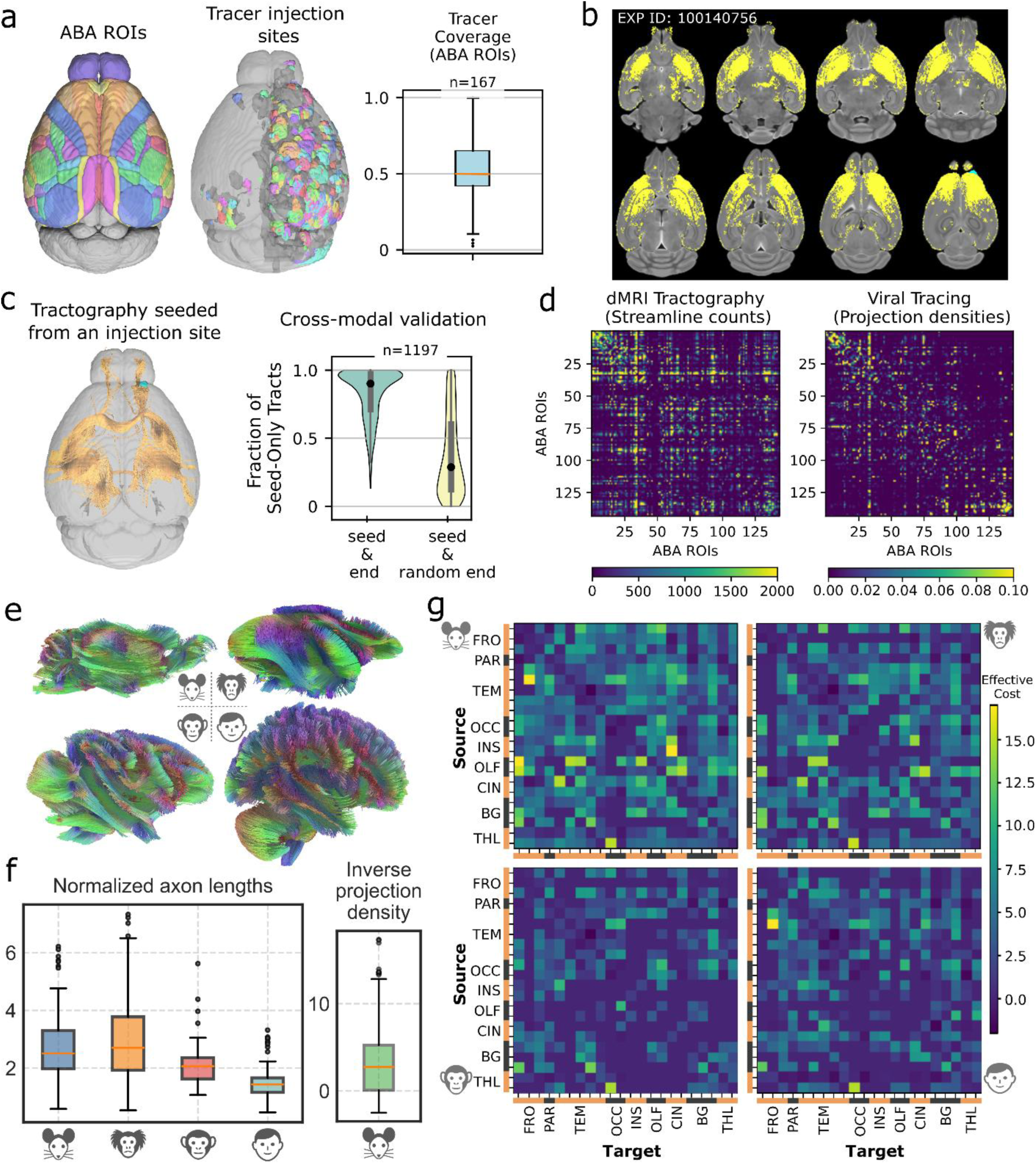
Construction and validation of directed connectomes across species. **(a)** Allen Brain Atlas (ABA) regions of interest (ROIs) with tracer injection sites (left) and distribution of ROI coverage by individual injections (right), showing ∼50% average coverage per region. **(b)** Representative viral tracer experiment (EXP ID: 100140756), with tracer-labeled projections overlaid in yellow on structural MRI slices. **(c)** Cross-modal validation of tracer–tractography correspondence. Tractography seeded from injection sites reproduced >90% of seed-only tracts when constrained with tracer-defined endpoints, far exceeding random controls (Wilcoxon signed-rank test, one-tailed, “greater”: W = 680,071, p < 0.0001, r = 0.80). **(d)** Connectivity matrices of diffusion MRI tractography (streamline counts, left) and viral tracing (projection densities, right) at the ABA ROI level, showing broadly consistent patterns of inter-regional connectivity. **(e)** Whole-brain diffusion MRI tractography reconstructions for mouse, marmoset, rhesus macaque, and human, illustrating species differences in scale and complexity of reconstructed pathways. **(f)** Distribution of normalized axonal lengths (left) and inverse projection densities (right), providing the inputs to the effective connection cost metric. **(g)** Directed connectivity matrices of effective connection cost for mouse, marmoset, rhesus macaque, and human, integrating tracer-defined polarity with species-specific axonal lengths. Lower costs correspond to shorter, stronger projections and form the basis for downstream path efficiency analyses.

Connectivity matrices at the ABA ROI level revealed broadly consistent organization between tractography-derived streamline counts and tracer-derived projection densities (**Fig. 2d**), supporting the integration of the two modalities. Whole-brain tractography^11^ across mouse, marmoset, rhesus macaque, and human highlighted striking species differences in scale and complexity of reconstructed pathways (**Fig. 2e**). To convert these data into directed connectomes, we combined tracer-derived polarity with species-specific axonal lengths, producing effective connection cost matrices (**Fig. 2g**). This biologically grounded metric reflects the physical and metabolic cost of axonal signaling by penalizing long, weak projections and favoring shorter, stronger ones. Distributions of normalized axonal lengths and inverse projection densities, shown in **Fig. 2f**, provide the inputs to this metric. In this formulation, shortest directed paths represent minimal-cost sequences of axonal projections and synaptic relays through which signals are most likely to exert strong influence on their targets.

### 2.2. Comparative path efficiency and cross-scale validation

We next asked how path efficiency differs across species. Real connectomes exhibited significantly lower minimum path costs than degree- and weight-preserving null networks with randomized tracer-defined directionality (**Fig. 3a**, left), demonstrating that polarity stabilizes network organization. When directionality was randomized, networks admitted inefficient detours and spurious shortcuts, inflating costs and producing extreme outliers (**Supplementary Fig. 1**).

**Figure 3.**
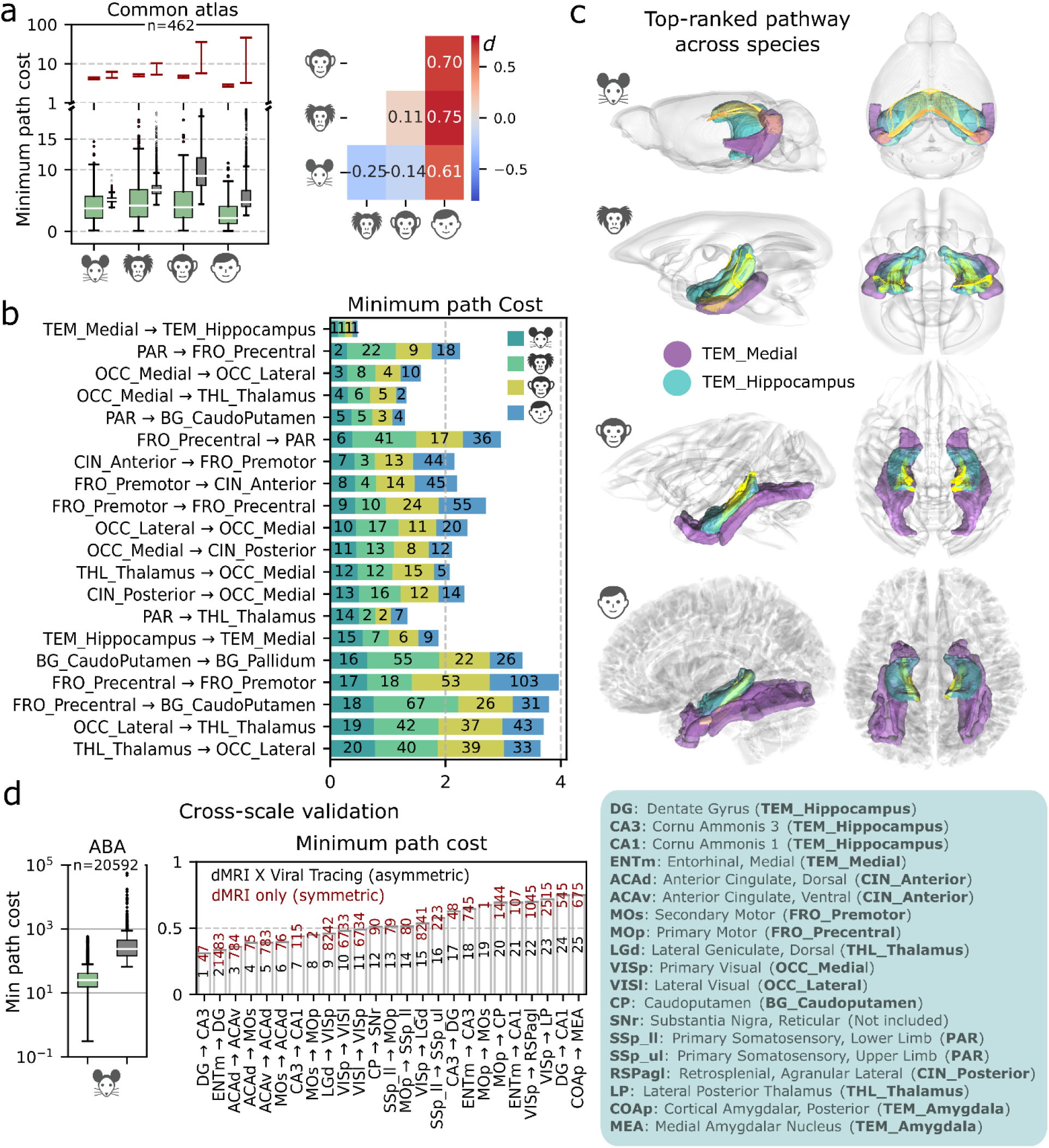
Comparative analysis of directed path efficiency across species with cross-scale validation. **(a)** Distribution of minimum path costs in directed connectomes derived from the common cross-species atlas. Real networks (green) are compared against 10,000 null networks generated by randomizing tracer-defined directionality (gray), showing that directionality stabilizes network organization and prevents inefficient detours. Right: pairwise interspecies comparisons summarized by Cohen’s *d*, with bootstrap resampling (n = 1,000) for statistical testing: Mouse vs. Marmoset *p* < 0.0001; Mouse vs. Rhesus *p* = 0.0410; Mouse vs. Human *p* < 0.0001; Marmoset vs. Rhesus *p* = 0.0840; Marmoset vs. Human *p* < 0.0001; Rhesus vs. Human *p* < 0.0001. **(b)** Top 20 most efficient mouse paths, sorted by cost, with corresponding values in other species shown as stacked colored bars. Conserved high-efficiency routes include medial temporal → hippocampus, thalamo–occipital, and precentral–parietal pathways. **(c)** Visualization of the top-ranked pathway across all species: medial temporal → hippocampus. 3D renderings highlight this conserved circuit in mouse, marmoset, rhesus macaque, and human. **(d)** Cross-scale validation within the Allen Brain Atlas (ABA). Left: distributions of minimum path costs for real networks (green) compared to 10,000 tracer-randomized nulls (gray), demonstrating results consistent with panel a. Right: bar plot of the 25 most efficient ABA-defined paths, with costs from dMRI-only symmetric connectomes shown in red for reference. Canonical hippocampal trisynaptic circuits (ENTm → DG, DG → CA3, CA3 → CA1), along with thalamo–occipital and sensorimotor loops, are recapitulated, confirming the biological fidelity of the framework at finer resolution. Together, these analyses demonstrate that tracer-informed directionality stabilizes network organization, recapitulates canonical hippocampal circuits, and enables comparative analysis of conserved versus lineage-specific efficiencies.

Bootstrap resampling of 1,000 iterations confirmed robust interspecies differences (**Fig. 3a**, right). Effect-size comparisons highlighted a pronounced human advantage: path costs were lower in humans relative to mouse (d = 0.61, p < 0.0001), marmoset (d = 0.75, p < 0.0001), and rhesus macaque (d = 0.70, p < 0.0001). Marmosets exhibited higher costs than both humans and rodents, likely reflecting constraints from longer axonal trajectories and less optimized routing in smaller primates. Rhesus macaques showed intermediate values (Mouse vs. Rhesus p = 0.0410; Marmoset vs. Rhesus p = 0.0840). This non-monotonic trajectory suggests that early primate brain expansion was associated with increased path costs, followed by a relative restoration of efficiency in higher primates through reallocation of network architecture.

Path-level analyses revealed both conserved and divergent influence channels. The medial temporal (ENTm) → hippocampus projection consistently ranked as the single most efficient pathway in all species (**Fig. 3b–c**), underscoring evolutionary preservation of memory-related circuitry^12,13^. Several other high-efficiency routes in the mouse, including thalamo–occipital (LGd ↔ VISp) and precentral–somatosensory (MOp ↔ SSp_ll) pathways, were also highly ranked in primates, reflecting broader conservation across cortical and subcortical systems.

At finer scale, analyses within the Allen Brain Atlas (ABA) provided a cross-scale validation of the framework in mouse (**Fig. 3d**). In addition to recovering the canonical hippocampal trisynaptic circuit— including the perforant path^14^ (ENTm → DG), mossy fibers^15^ (DG → CA3), and Schaffer collaterals^16^ (CA3 → CA1)—the ABA analysis also refined macro-scale connections represented in the common atlas, such as reciprocal LGd ↔ VISp interactions (thalamo–occipital) and Mop ↔ SSp_ll pathways (precentral–somatosensory). Distributions of minimum path costs showed results consistent with those at the macro scale, with real networks outperforming 10,000 direction-randomized nulls. Bar plots of the top 25 ABA-defined paths further demonstrated that symmetric dMRI-only connectomes failed to capture canonical organization (shown in red), whereas tracer-informed networks aligned with established anatomy.

Together, these results show that integration of tracer-derived directionality with species-specific tractography recapitulates the network’s most influential channels across scales. Even when applied to a coarser cross-species atlas, the framework remained consistent with finer pathways defined in the ABA, confirming the biological fidelity of the integration approach for comparative analysis.

### 2.3. Regional, role-specific cross-species differences in path efficiency

Having established whole-network efficiency profiles and their cross-scale validation, we next asked where efficiency reallocations were concentrated at the level of specific regions and roles. For each region, paths were grouped by whether it served as a source (outflow) or a target (inflow), and efficiency distributions were compared across species. Six region–role combinations showed significant cross-species differences after Kruskal–Wallis tests with FDR correction (**Table 1**).

**Table 1.** Regional-Level Cross-Species Comparisons Using Kruskal–Wallis Tests.

| Common Regions | As Source |  | As Target |  |
| --- | --- | --- | --- | --- |
|  | Diff | P | Diff | P |
| FRO_Precentral (PreC) | 3.49 | 0.788 | 1.33 | 0.998 |
| FRO_Premotor (PreM) | 0.74 | 0.998 | 2.53 | 0.899 |
| FRO_Prefrontal (Pref) | 0.80 | 0.998 | 0.33 | 0.998 |
| PAR | 0.22 | 0.998 | 0.54 | 0.998 |
| TEM_Superior (TemS) | **12.41** | 0.045 | 4.85 | 0.537 |
| TEM_Inferior (TemI) | **14.28** | 0.022 | 7.63 | 0.199 |
| TEM_Medial (TemM) | 0.90 | 0.998 | 0.35 | 0.998 |
| TEM_Hippocampus (Hipp) | 3.73 | 0.788 | 0.51 | 0.998 |
| TEM_Amygdala (Amyg) | 5.59 | 0.419 | 1.40 | 0.998 |
| OCC_Lateral (OccL) | 0.18 | 0.998 | 0.10 | 0.998 |
| OCC_Medial (OccM) | 0.36 | 0.998 | 0.08 | 0.998 |
| INS_Anterior (InsA) | **34.99** | 0.000 | **26.72** | 0.000 |
| INS_Posterior (InsP) | 5.64 | 0.419 | 0.73 | 0.998 |
| OLF_Anterior (OlfA) | **18.10** | 0.005 | 9.80 | 0.089 |
| OLF_Piriform (Pir) | 11.63 | 0.055 | **30.94** | 0.000 |
| CIN_Anterior (CingA) | 3.35 | 0.788 | 1.97 | 0.998 |
| CIN_Posterior (CingP) | 0.04 | 0.998 | 0.24 | 0.998 |
| BG_CaudoPutamen (CPu) | 3.40 | 0.788 | 0.86 | 0.998 |
| BG_Accumbens (NAc) | 8.93 | 0.121 | 9.91 | 0.089 |
| BG_Pallidum (Pall) | 2.74 | 0.899 | 2.57 | 0.899 |
| THL_Thalamus (Thal) | 0.72 | 0.998 | 0.23 | 0.998 |
| THL_Hypothalamus (Hypo) | 10.47 | 0.082 | 2.53 | 0.899 |

The most consistent human gains occurred in the anterior insula (source and target) and in superior temporal cortex (source), all showing elevated path efficiencies relative to other species (**Fig. 4a**). Regional maps of divergence further highlighted these loci, along with inferior temporal and anterior olfactory cortex for source roles, and anterior insula and piriform cortex for target roles (**Fig. 4b**). Pairwise species comparisons using Dunn’s post hoc tests (Supplementary Table 2) confirmed that human advantages were most pronounced in anterior insula (both roles) and superior temporal (source), while marmoset retained the strongest olfactory influence in both outflows and inflows (**Fig. 4c**).

**Figure 4.**
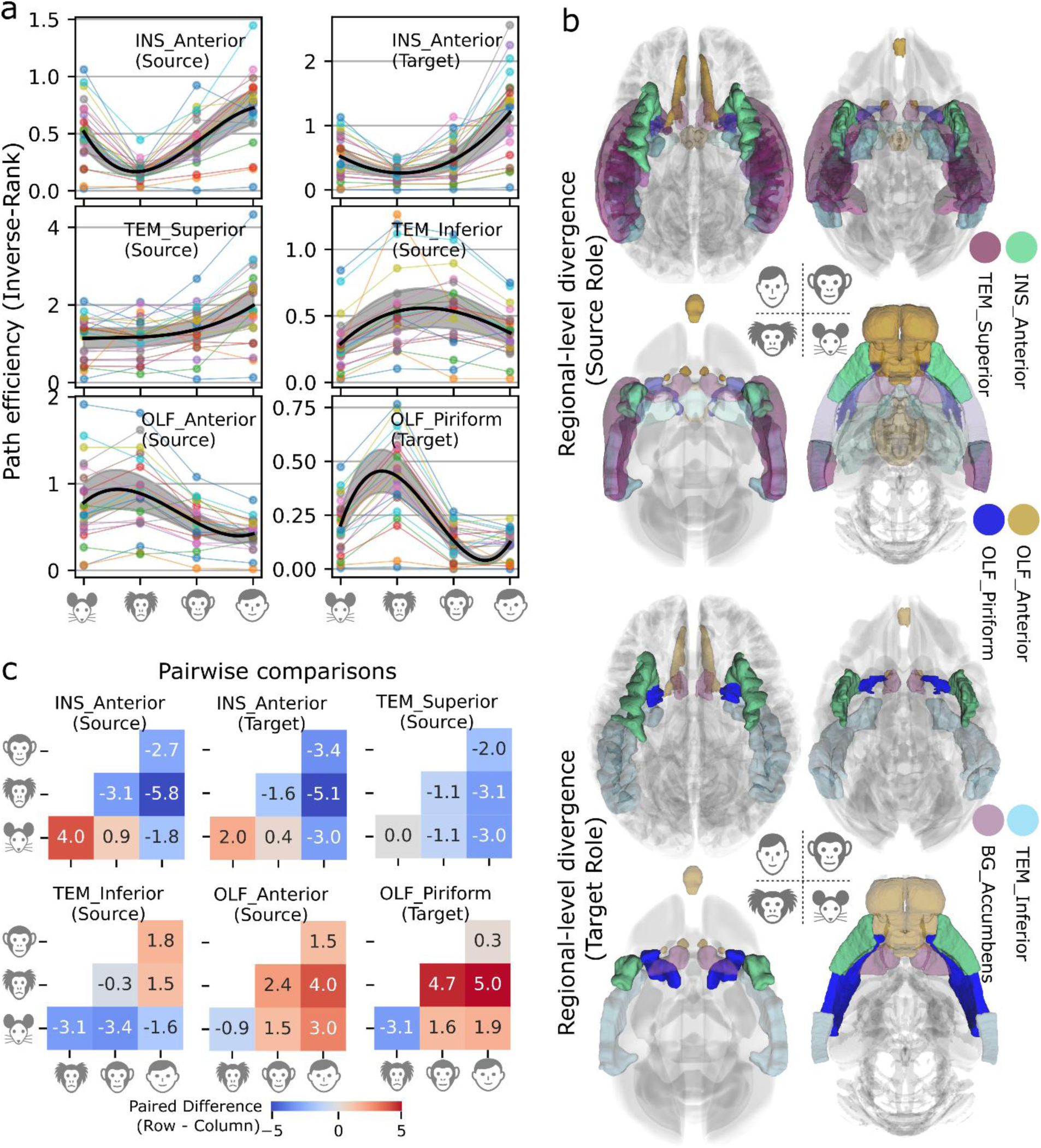
Regional and role-specific reallocations of path efficiency across species. **(a)** Path efficiency distributions (inverse-rank values) for the six region–role combinations showing significant cross-species differences by Kruskal–Wallis tests (see Table 1). Thin colored lines represent individual paths, thick black curves indicate smoothed species means, and shaded bands denote 95% confidence intervals. Human gains are most evident in anterior insula (source and target) and superior temporal (source), rhesus maxima appear in inferior temporal (source), and marmoset maxima occur in olfactory inflows and outflows. **(b)** Regional maps of cross-species divergence, with Table 1 Kruskal–Wallis differences (Diff values) projected onto anatomical ROIs. Regions with Diff > 5.0 are highlighted. Source-role analyses identify the anterior insula, superior temporal, inferior temporal, and anterior olfactory cortex as loci with significant divergence in at least one species. Target-role analyses identify the anterior insula and piriform cortex as regions with the strongest species differences. **(c)** Pairwise species comparisons of path efficiency for the six significant region–role combinations, using Dunn’s post hoc tests (see Supplementary Table 2). Heatmaps show mean paired differences (row – column); blue indicates higher efficiency in the column species, red indicates higher efficiency in the row species. The clearest human advantages occur in anterior insula (both roles) and superior temporal (source), while marmoset retains high olfactory influence (anterior source and piriform target). Together, these analyses pinpoint the specific cortical and subcortical loci where efficiency has been differentially reallocated across lineages, highlighting human amplification of temporal–insula roles, primate specialization of inferior temporal outflows, and marmoset preservation of olfactory pathways.

Inferior temporal outflows tended to be stronger in primates than in mouse, with rhesus macaques reaching the highest values, followed by marmosets and humans. These patterns suggest a primate-specific enhancement of ventral visual stream pathways^17–19^ that peaked in rhesus, while humans may have shifted efficiency toward insula-related interactions. In contrast, olfactory roles showed the opposite trajectory, with strongest efficiencies in marmoset and weakest in humans. Anterior olfactory outflows to piriform and temporal regions were particularly enhanced in marmoset relative to both rodents and higher primates, consistent with chemosensory-based communication and ecological adaptation^20,21^.

### 2.4. Partner-specific contributions to regional role differences

To clarify which circuits underlie these regional reallocations, we constructed partner-level Δ-connectomes for five representative region–role combinations with the strongest cross-species divergence (**Fig. 5a**). Each focal region was paired with its top five partner nodes, ranked by absolute mean Δ values across pairwise comparisons. These analyses identified the partner pathways that drive regional differences in efficiency.

**Figure 5.**
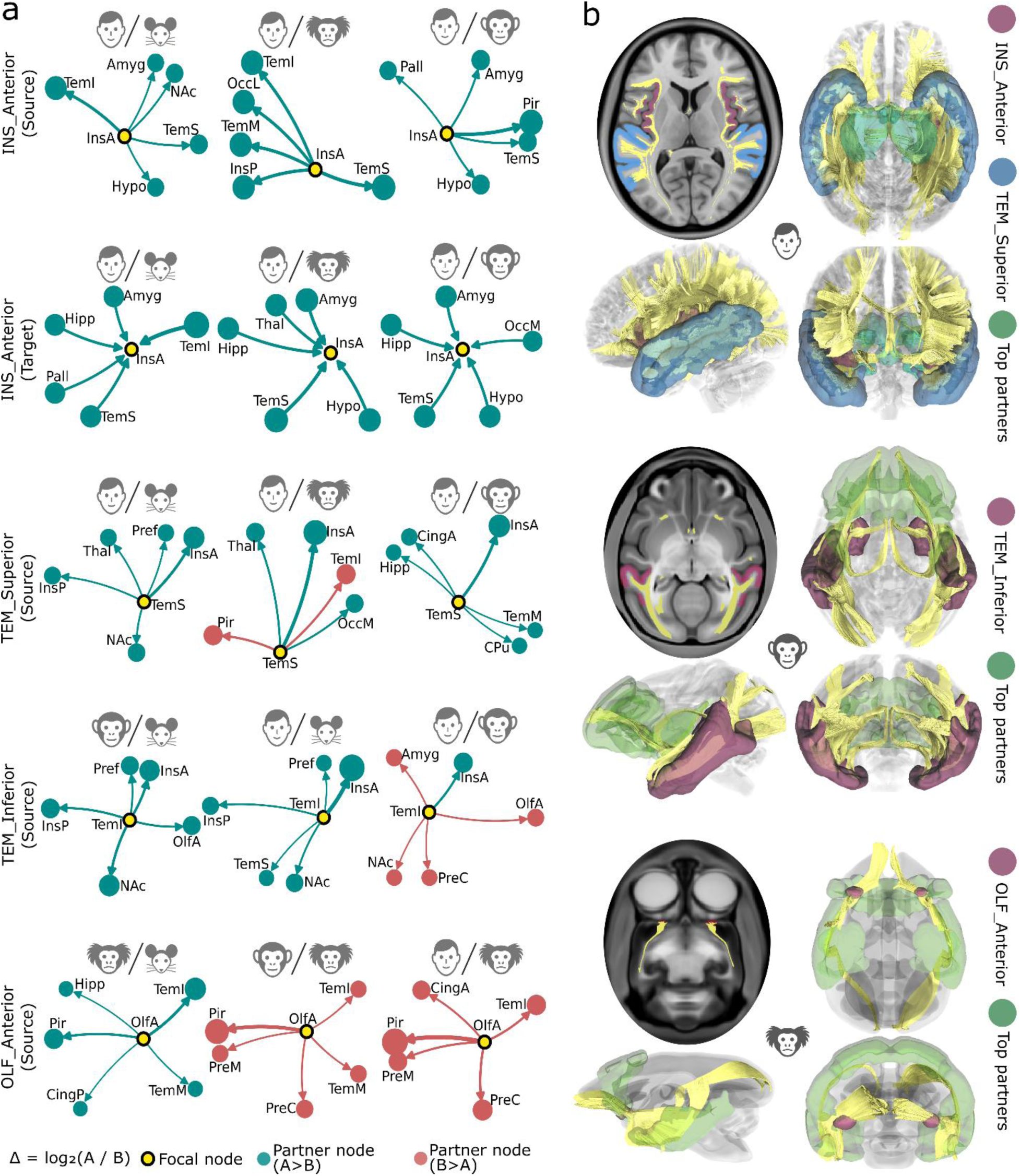
Partner-specific contributions to regional role differences. **(a)** Partner-level Δ-connectomes for five representative region–role combinations with significant cross-species divergence: anterior insula (source, target), superior temporal (source), inferior temporal (source), and anterior olfactory (source). Each subgraph shows the focal region (yellow) and its top five partner nodes, ranked by absolute mean Δ values across pairwise species comparisons. Node color indicates the direction of difference (green = higher in the numerator species, red = higher in the denominator species), node size and edge width scale with |Δ|, and edge direction reflects the focal role (outflow or inflow). These graphs identify specific partner pathways driving species differences, including strengthened temporal–insula reciprocity in humans, ventral-stream outflows in rhesus, and olfactory outflows in marmosets. **(b)** Brain renderings of focal regions and their partner pathways, grouped by the species that showed the maximum efficiency for each region–role combination. Top: human maxima in anterior insula and superior temporal cortex. Middle: rhesus maxima in inferior temporal cortex. Bottom: marmoset maxima in anterior olfactory cortex. Focal regions and partners are color-coded within each species rendering, while yellow overlays depict white-matter tracts linking the focal region to its partners. Together, these analyses reveal the partner-specific basis of regional role differences, pinpointing strengthened temporal–insula circuits in humans, ventral visual stream specialization in Old World primates, and preserved olfactory routes in New World primates.

In humans, anterior insula and superior temporal regions both showed strengthened reciprocal connections with each other and with prefrontal partners. Prefrontal participation was most evident in human–mouse contrasts, whereas human–primate comparisons emphasized enhanced temporal–insula reciprocity. In rhesus macaques, inferior temporal outflows targeted prefrontal and Insula regions as well as nucleus accumbens, consistent with ventral visual stream specializations and integration with subcortical targets^17–19^. In marmosets, anterior olfactory outflows to piriform, temporal, and frontal regions emerged as the dominant signature, reinforcing the preservation of olfactory influence in New World primates.

Three-dimensional brain renderings illustrated these focal regions and their top partner pathways, grouped by the species that showed the maximum efficiency for each role (**Fig. 5b**). Human maxima were observed in anterior insula and superior temporal cortex, rhesus maxima in inferior temporal cortex, and marmoset maxima in anterior olfactory cortex. These renderings highlight the anatomical circuits underlying lineage-specific reallocations: strengthened temporal–insula interactions with prefrontal contributions in humans, inferior temporal outflows to cortical and subcortical partners including NAc in rhesus macaques and preserved olfactory routes in marmosets.

Taken together, the regional and partner-level analyses reveal three domain-specific evolutionary trajectories. First, humans show amplification of temporal–insula–frontal pathways, with anterior insula and superior temporal regions jointly strengthened and prefrontal participation most evident in human–mouse contrasts, consistent with enhanced multimodal integration and language-related functions^22–24^. Second, primates exhibit an expansion of ventral visual stream pathways^25,26^, peaking in rhesus macaques where inferior temporal outflows extend to prefrontal, superior temporal, and subcortical partners including nucleus accumbens. Third, marmosets maintain relatively strong olfactory influence, expressed through anterior olfactory and piriform pathways, in line with chemosensory communication and ecological specialization^20,21^. These lineage-specific reallocations indicate that path efficiency has been differentially distributed across species, producing both human-associated gains and primate- or marmoset-specific specializations.

## 3. Discussion

### 3.1. Viral Tracer Integration and Cross-Species Directed Connectomes

A central challenge in comparative connectomics is that viral tract-tracing, the gold standard for determining projection directionality, remains practically and ethically restricted to mice and a few other small-brain species. Its invasive nature makes comprehensive application in humans or great apes impossible, leaving non-invasive diffusion MRI as the only feasible modality for large-brain primates. Diffusion tractography, however, is inherently symmetric: reconstructed streamlines yield bidirectional connectivity estimates, obscuring the unidirectional propagation of action potentials. This reciprocity limits the biological interpretability of dMRI-derived networks, especially in frameworks such as graph theory or dynamical systems modeling where directionality is essential^3,6^.

Against this backdrop, our framework leverages the uniquely comprehensive viral tracing dataset in mouse (∼1,200 injection experiments) as a directional scaffold, combined with species-specific dMRI-derived axonal lengths. This hybrid approach exploits the complementary strengths of both modalities: empirically validated projection asymmetries from tracing and whole-brain coverage from tractography. As shown in **Fig. 3a** and **Supplementary Fig. 1**, incorporating directionality stabilizes network organization across species. Randomizing tracer-defined directions causes networks to admit implausible detours and spurious shortcuts, generating extreme outliers and inflated path costs. By contrast, preserving true directionalities yields efficient and narrowly distributed cost profiles, even when paired with the wiring lengths of different species. These findings suggest that mouse-derived directionality provides a biologically meaningful constraint—one that grounds efficiency analyses across species, and without which the organizational structure would likely be severely disrupted.

Although based on mouse viral tracing, this integration provides precisely the directional constraint needed to uncover species-specific adaptations relative to a shared scaffold. What is introduced from the mouse is not its entire connectivity architecture, but only the empirically validated polarity of projections: the direction in which signals flow between regions. The spatial organization of white-matter trajectories, i.e. their lengths and embedding within each brain, is derived entirely from species-specific diffusion MRI, allowing us to examine how wiring constraints may have been shaped differently across lineages. By combining projection polarity from mouse tracing with species-specific axonal lengths, the framework makes it possible to reveal lineage-specific trade-offs: humans showed elevated efficiencies in long association pathways forming a temporal–insula–frontal circuit. This circuit specifically involved superior temporal, anterior insula, and prefrontal cortices, providing a structural substrate for multimodal integration and language-related functions. Rhesus macaques showed relative peaks in ventral visual outputs, and marmosets retained prominent olfactory–piriform influence (**Figs. 4&5**). In this way, directionality provides the missing constraint for path search, while each species’ white-matter organization shapes a distinct efficiency landscape.

At present, there is no comprehensive alternative for defining directed human connectomes. Functional approaches such as dynamic causal modeling^5,27–29^ have been widely applied to fMRI to infer directed connectivity, but their interpretation remains constrained by hemodynamic variability^30,31^ across cortical regions. Differences in vascular dynamics introduce uncertainty in the temporal alignment of regional activation patterns, making causal interactions difficult to assess with rigor. Furthermore, causal relationships inferred from hemodynamic signals may not directly correspond to neuronal-level propagation^32–34^, underscoring the need for structurally informed models that bridge scales. By grounding our networks in empirical structural asymmetries, we provide a biologically validated scaffold for directionality that avoids these pitfalls.

Importantly, the directed shortest paths identified in our models can be interpreted as routes of forward action potential propagation—minimal-cost sequences of axonal projections and synaptic relays that maximize downstream impact per unit wiring investment. This framing bridges scales: the directed connectomes derived here are not statistical abstractions, but plausible mechanistic blueprints for how neuronal signals traverse large-scale networks. In turn, they provide a foundation for cross-species mechanistic modeling of cognition with spiking neural networks, where biologically grounded directional paths are essential for simulating signal flow, coherence, and emergent dynamics.

Together, these findings suggest that polarity derived from mouse viral tracing provides a biologically meaningful constraint that stabilizes network organization across species. By preventing implausible detours and anchoring path search to realistic signaling directions, the framework yields networks that are both efficient and more consistent with canonical circuits. These results support the technical validity of integrating tracer polarity with species-specific tractography. In the following section, we consider more broadly how mouse tracer data can be used as a directional scaffold for cross-species connectomics, and the conceptual and translational implications of generalizing polarity across species.

### 3.2. Mouse Tracer Data as a Directional Scaffold

The integration presented here depends on the uniquely comprehensive viral tracing dataset in the mouse, consisting of ∼1,200 anterograde injections compiled by the Allen Institute. A valid concern is whether projection polarity observed in the mouse can be generalized to larger-brained primates and humans. Our framework does not assume that the mouse connectome as a whole is conserved across species. Rather, it transfers only the empirically validated polarity of projections, the direction in which signals travel between regions, while leaving the topology and wiring constraints entirely species-specific through dMRI tractography.

Several lines of evidence support this design choice. Many canonical hippocampal circuits, such as the entorhinal–hippocampal pathway and the trisynaptic loop, are well established as conserved across mammals. Place cell and grid cell dynamics, originally discovered in rodents, have also been demonstrated in the human hippocampal formation^35–37^, underscoring deep conservation of hippocampal circuitry across species. Our own results reinforce this: introducing polarity stabilizes network organization across species **(Fig. 3a**), and integration of tracer with tractography recapitulates the trisynaptic circuit, whereas dMRI-only symmetric connectomes fail to capture it (**Fig. 3d**). Taken together, these findings highlight that polarity is not arbitrary but a biologically grounded constraint that improves both network stability and fidelity to canonical circuits.

It is also important to recognize that most current human network neuroscience relies on undirected connectomes, which necessarily limit the interpretation of neuronal signal flow^3,5^. By incorporating polarity from invasive experiments, we move toward more mechanistic accounts of network function. More broadly, mouse models have been indispensable in advancing understanding of mammalian neural circuit function, from canonical circuits^38,39^ to large-scale atlases^4,40,41^, with disease models^42,43^ providing additional translational relevance. Substantial resources and animal lives have been invested to generate invasive datasets^4,44^, and it is both scientifically and ethically important to make responsible use of them, even while acknowledging their translational uncertainty. To do so responsibly, these datasets must be leveraged with care: our cross-species framework makes use of polarity only at a macro-scale level, where divergences are minimized. For example, we deliberately keep the parietal region (PAR) collapsed into a single division across species to account for differences in its finer subdivisions^7^. Such macro-level analysis, combined with species-specific tractography, mitigates many concerns about overgeneralization. Moreover, even if future primate tracer resources become widely available, uncertainty in cross-species alignment will inevitably remain. Nevertheless, our framework provides a foundation for systematically incorporating such future data while maintaining consistency across scales.

Beyond comparative mapping, these directed connectomes also provide a foundation for mechanistic models of large-scale brain dynamics^45^. By anchoring shortest paths to empirically validated polarity, they yield plausible blueprints for how activity may propagate through real circuits, which can in turn constrain simulations of signal flow, coherence, and emergent network behavior. In this sense, the ethical and scientific value of mouse tracing extends beyond comparative anatomy: it supplies the directional constraints needed to bridge invasive data with noninvasive imaging, and to link structure with dynamics in a way that undirected connectomes cannot.

### 3.3. Cross-Species Comparisons

A consistent finding across species was the high efficiency of the entorhinal–hippocampal pathway, which remained top-ranked in all connectomes (**Fig. 3b**). This preservation underscores how certain core circuits have been maintained through evolution, even as other systems appear to have been differentially emphasized. In contrast, lineage-specific reallocations of efficiency were also evident. For example, olfactory influence peaks in marmosets rather than mice, reflecting the strong chemosensory investment of New World primates and the relatively weaker association systems that may have allowed olfactory routes higher in the efficiency hierarchy. Rodents, by comparison, emphasize somatosensory systems, which may have reduced the relative prominence of olfactory pathways despite their absolute strength.

These contrasts highlight why evolutionary change is best interpreted in terms of relative prioritization rather than absolute wiring costs. Absolute path costs (axon length × inverse projection strength) can be size-corrected to enable interspecies comparison, but they primarily index the physical and metabolic expense of individual routes. In contrast, our rank-based efficiency situates each connectome within its own competitive landscape: shortest directed paths are ranked against all alternatives, so differences reflect shifts in network prioritization rather than absolute expense. This framing exposes trade-offs that absolute costs obscure: in humans, temporal–insula–frontal routes rise to the top, with prefrontal contributions most evident in human–mouse contrasts and temporal–insula reciprocity dominating in human–primate comparisons, while olfactory routes fall to the bottom. In marmosets, weaker association systems allow olfactory paths to occupy higher ranks. In rhesus macaques, inferior temporal outflows peak toward prefrontal and insular cortices and extend to nucleus accumbens, reflecting ventral-stream specializations without further human amplification. By integrating directionality with whole-network path search, the ranking captures how evolutionary pressures may have constrained influence hierarchies, identifying which circuits tend to be promoted or demoted, rather than only how long or strong individual edges are.

Taken together, our findings suggest that evolution reshaped cortical communication not through uniform scaling, but by reallocating efficiency across a balance of conserved core pathways and lineage-specific specializations. Humans showed elevated efficiency in a temporal–insula–frontal circuit, potentially supporting the integrative capacities underlying complex cognition and language^22–24^. Old World primates such as rhesus macaques exhibited high efficiency in inferior temporal outflows directed toward prefrontal and insular cortices and to nucleus accumbens, consistent with ventral-stream circuits for object and face recognition^25,26^ that also incorporate subcortical integration. New World primates such as marmosets retained relatively strong olfactory influence, consistent with their reliance on chemosensory cues for social communication and ecological adaptation^20,21^. Alongside these divergent reallocations, conserved memory-related canonical projections, the entorhinal–hippocampal pathway^12,13^ persisted across all species, anchoring the efficiency landscape. Together, these results indicate that cortical networks may have diversified through lineage-specific peaks built alongside shared foundations, with humans, rhesus, and marmosets each showing distinct patterns in the trade-offs imposed by expanding brain systems. These interpretations follow from the foundational assumption of comparative neuroscience: that cross-species differences reflect evolutionary divergence, and that statistical contrasts reveal the selective pressures shaping neural organization across lineages^9^.

### 3.4. Limitations and Future Directions

Several limitations of this work should be acknowledged. First, projection polarity was derived exclusively from viral tracing in the mouse. This issue and its implications are discussed in detail above (see Discussion: Mouse Tracer Data as a Directional Scaffold). Briefly, while the dataset is uniquely comprehensive, some asymmetries may not be conserved across species, and future efforts incorporating primate tracer resources or new experimental modalities will be needed to refine the approach. Second, resolution differences between the common atlas and finer species-specific atlases impose constraints. Some structures, such as the claustrum or subthalamic nucleus, could not be reliably included^7^, and the coarse common atlas resolution may obscure subfield-level specializations within regions like hippocampus or thalamus. Third, path efficiency emphasizes routing through shortest directed paths, but real neural dynamics can exploit redundant or parallel routes; complementary modeling approaches may provide a fuller account of interregional communication strategies.

Looking ahead, the integration of tracer-derived directionality with tractography offers several promising extensions. Higher-resolution parcellations, especially at the subnuclear level in limbic and thalamic systems, will enable more detailed analyses of intra-system organization. Additional cross-species tracer datasets, including ongoing work in marmoset and macaque, may allow polarity constraints to be derived directly within primates. Finally, coupling the directed connectomes with computational models, ranging from network control theory to spiking neural networks, could link the structural efficiency patterns observed here to functional dynamics and behavioral capacities.

## 4. Materials and Methods

### 4.1. Diffusion MRI Data and Processing

Diffusion MRI templates were constructed for human, rhesus macaque, marmoset, and mouse, each based on group-averaged datasets with species-appropriate acquisition and reconstruction parameters. For the human template, we used a total of 1,065 healthy adult scans acquired with a multi-shell diffusion scheme (b-values = 990, 1985, and 2980 s/mm²; 90 directions per shell)^46^. The in-plane resolution and slice thickness were 1.25 mm. Diffusion data were reconstructed using q-space diffeomorphic reconstruction^47^, with restricted diffusion quantified using a model-free method^48^. Deterministic white matter fiber tracking^49^ was applied, followed by trajectory-based tract recognition^50^ and topology-informed pruning^51^ to minimize false positives.

For the rhesus macaque template, we used 44 scans acquired with a multi-shell diffusion scheme (b-values = 500, 1000, and 2000 s/mm²; 6, 30, and 30 directions, respectively). Images were acquired at 1 mm isotropic resolution and resampled to 0.5 mm. Susceptibility distortions were corrected using TOPUP (Tiny FSL package, a re-compiled version of FSL TOPUP)^52^, and b-tables were quality-checked by an automated routine^53^. Data were reconstructed in MNI space using generalized q-sampling imaging^54^, with restricted diffusion quantified as above. Deterministic fiber tracking^49^ with augmented tracking strategies^50^ was employed to improve reproducibility.

For the marmoset template, we used 126 scans acquired with a multi-shell diffusion scheme (b-values = 1013 and 3019 s/mm²; 30 and 60 directions, respectively). The in-plane resolution was 0.35 mm with a slice thickness of 0.7 mm, and images were resampled to 0.20 mm isotropic. Diffusion data were reconstructed in MNI space using q-space diffeomorphic reconstruction^47^ to obtain spin distribution functions^54^, with a diffusion sampling length ratio of 1.25 and an output resolution of 0.20 mm isotropic. Restricted diffusion was quantified using restricted diffusion imaging^48^.

For the mouse template, we used 22 C57BL/6 brains acquired at ultra-high resolution (0.025 mm isotropic) with a single-shell HARDI sequence (b = 3000 s/mm²; 114 directions). Diffusion data were reconstructed in MNI space using q-space diffeomorphic reconstruction^47^ to obtain spin distribution functions^54^, with an output resolution of 0.025 mm isotropic. Tractography was performed using deterministic fiber tracking^49^ with augmented tracking strategies^50^.

All templates were aligned to standard stereotaxic coordinate systems, reconstructed using consistent model-free approaches, and processed with deterministic tractography, ensuring comparability across species.

### 4.2. Mouse Viral Tracer Integration and Directed Path Modeling

#### 4.2.1. Viral Tracer Data and Preprocessing

We used a publicly available mouse mesoscale connectome dataset^4^ comprising 1,199 anterograde viral tracing injection experiments reported in the Allen Mouse Brain Connectivity Atlas database. For each experiment, both injection density and projection density volumes were downloaded in NRRD format via the Allen Brain Atlas API (http://api.brain-map.org/), converted to NIfTI, and spatially registered to the mouse diffusion MRI space using rigid and nonlinear transformations. Two experiments contained no labeled voxels and were excluded from analysis, leaving 1,197 injections for computation. Volumes were binarized to remove floating-point artifacts, yielding binary masks of injection and projection coverage. All tracer and tractography data were mapped to a previously developed common hierarchical atlas^7^ that defines 22 homologous cortical and subcortical regions across species. This ensured one-to-one correspondence of ROIs for integration and comparison.

#### 4.2.2. Estimation of Directed Projection Strengths

Let *A* be the atlas label volume (*0* = background; *1…R* = region IDs), *S*_*k*_ the binarized injection mask for experiment *k*, and *P*_*k*_ the binarized projection mask for the same experiment. The indicator function 1[.] equals 1 if the condition holds and 0 otherwise.

Injection fraction for region *i* (experiment *k*):

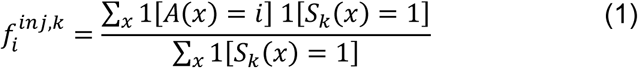

Here 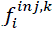 denotes the fractional coverage of source region *i* by experiment *k*; experiments with partial injections contribute proportionally to their coverage, while zero-coverage cases contribute nothing. This ensures that directed strengths are weighted by the effective fraction of each region covered by the injection. Projection fraction for region *j* (experiment *k*):

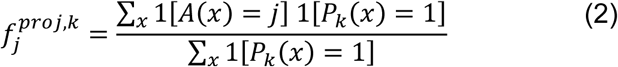

Experiment-level weight from *i* → *j*:

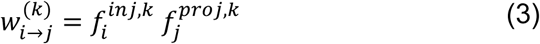

Final directed projection matrix (sum over valid experiments; *N* = 1197):

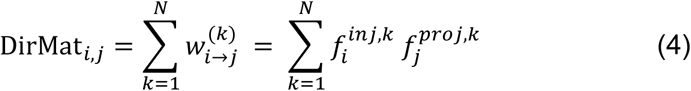

This procedure allocates each experiment’s injected signal across source regions 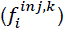 and its projection signal across target regions 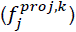, then form a sum of outer products across experiments to estimate directed strengths *i* → *j* in atlas space. Background voxels (*A* = 0) are excluded from all sums. Thus, the final directed matrix represents a coverage-weighted sum across all experiments, reducing bias from incomplete tracer injections.

For the fine-scale mouse analysis, the same computation is performed in Allen Brain Atlas (ABA) space. For the cross-species analysis, the directed strengths computed in the mouse MRI atlas space (common atlas version) are subsequently integrated with species-specific diffusion MRI connectivity to derive *effective connection costs* and *path efficiencies*.

#### 4.2.3. Diffusion MRI Connectivity

For each species (mouse, marmoset, rhesus macaque, human), diffusion MRI tractography was performed to generate a species-specific undirected connectivity matrix of mean axonal lengths^11^ *L*_*ij*_ = *L*_*ji*_ between all common atlas region pairs. Mean lengths were expressed in millimeters and computed only for region pairs with a nonzero number of streamlines. Tractography was performed using whole brain seeding. For the mouse brain, the tractography output was mapped to the same diffusion MRI atlas space used for registering the viral tracer data, ensuring voxel-level correspondence for subsequent multimodal integration.

#### 4.2.4. Integration of Modalities into Effective Connection Cost Matrices

Mouse-derived directed projection strengths *DirMat_i,j_* in common atlas space were applied to the dMRI-derived axonal lengths *L*_*ij*_ for each species to compute the *effective connection cost*:

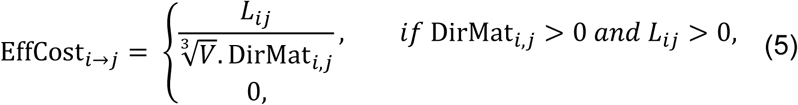

where *V* is the total white matter volume of the species (in *mm*^3^), used to normalize for brain size. A connection was defined only if both modalities supported it (nonzero DirMat_*i*,*j*_and nonzero *L*_*ij*_), reducing false positives from spurious tractography streamlines or tracer noise. Although axonal length is symmetric, EffCost_*i*→*j*_ ≠ EffCost_*j*→*i*_ because DirMat_*i*,*j*_ encodes directional projection strength.

Effective connection cost reflects the physical and metabolic investment required per unit of potential influence between two regions. For connections of equal length, a stronger projection yields a lower effective cost because it offers more potential influence per unit wiring investment. In contrast, weak projections of the same length are metabolically more wasteful, sustaining the same infrastructure for reduced influence yield.

Effective connection cost matrices were computed between homologous common-atlas ROIs across species. This set reflects specific consolidations necessitated by species-specific anatomical resolution: (i) Because the mouse atlas lacks parietal lobe subdivisions, we retained an undivided PAR region for all species (Level-1 definition for Parietal Lobe), (ii) In primates, the caudate and putamen were merged into a single BG_CaudoPutamen ROI to match the undivided form in rodents. These ROI definitions follow the constraints of the common atlas, ensuring consistency across species. With these constraints, the number of valid directed paths is 22×21=462. These same ROI definitions were used in all downstream path efficiency analyses, including the path-level ranking and regional-level comparisons.

#### 4.2.5. Path Efficiency Analysis

Minimum path costs between all region pairs were computed from the effective connection cost network as the lowest cumulative cost along any valid directed route, using a shortest directed path algorithm^8,55^.

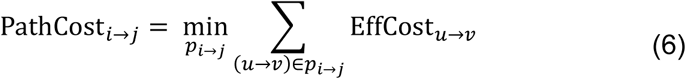

where *p*_*i*→*j*_ is any directed path (sequence of directed edges) from region *i* to *j*. Only edges with nonzero mapped projection strengths (DirMat_*u,v*_ > 0) were considered valid in the path search.

Wiring minimization alone may lack the explanatory power to fully account for the complex topology of the brain connectome^56^. In the current study, the minimization function is multi-objective, combining axonal length and the inverse of projection density. This favors shorter axons with stronger projections, yielding directed shortest paths that are simultaneously length-minimizing and projection-strength–maximizing.

Biologically, these minimal-cost directed paths approximate the forward propagation of action potentials from source to target through the most efficient influence channels. They represent sequences of axonal projections and synaptic relays that minimize wiring investment while maximizing potential impact, providing a mechanistic basis for modeling large-scale cortical dynamics.

#### 4.2.6. dMRI-only (symmetric) connectomes

To generate a dMRI-only version of the effective connection cost, we substituted the asymmetric projection density matrix (DirMat) in Equation 5 with the symmetric tractography matrix, where edge weights reflected inter-regional streamline density. This yielded an undirected cost formulation in which axonal length was combined with symmetric connectivity estimates, without incorporating tracer-derived polarity.

#### 4.2.7. Null models for path efficiency

For statistical comparison, we generated null distributions of minimal path cost using two complementary models, both based on row-wise permutation of nonzero entries within each connectivity matrix. In the tracer-randomized null, polarity assignments were shuffled across outgoing edges while preserving the number of nonzero outflows per node, thereby maintaining degree distribution but disrupting biological directionality. In the dMRI-randomized null, the same procedure was applied to the tractography matrix while tracer-defined polarity was kept intact. For each species, 10,000 randomized networks were generated to estimate null distributions of minimal path cost, with dMRI-only shuffle results reported in the Supplementary Information.

### 4.3. Cross-species Comparisons

#### 4.3.1. Cross-species Path Efficiency Ranking

To compare path efficiency across species, we ranked all valid inter-regional paths by their minimum effective path cost (Equation 6), with lower-cost paths receiving higher ranks. We then computed an inverse-rank efficiency score, 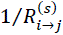, for species *s*, providing a normalized measure that allows comparison across species with different brain sizes and network scales. Since most paths have relatively low efficiency and the score distribution is right-skewed, a monotonic log transformation was applied for visualization purposes to expand the mid-range values and improve clarity in plots. However, this transformation was not used in subsequent Δ-connectome calculations.

#### 4.3.2. Regional-Level Statistical Comparisons

While all valid directed paths (n=462) were assigned efficiency ranks (see Methods, Path efficiency analysis), formal cross-species comparisons were performed at the regional level, treating each region separately in its role as a source (outgoing edges) or target (incoming edges). These analyses used the same 22 homologous ROIs defined for the effective connection cost matrices, ensuring one-to-one correspondence across species.

For a given region–role combination, we extracted the efficiency values of all valid paths in which that region served in the specified role, yielding one distribution per species. We then applied a Kruskal–Wallis test to assess whether distributions differed among species. When significant, we performed pairwise post hoc Dunn’s tests with false discovery rate (FDR) correction.

This approach reduces the number of statistical tests by evaluating distributions of all paths linked to a region rather than testing each path independently. While low-efficiency connections still contribute to the analysis, they are interpreted in aggregate alongside higher-efficiency connections from the same region, shifting the emphasis from isolated edges to anatomically coherent loci of potential divergence. This prioritizes direct anatomical interpretability and focuses on evolutionarily meaningful patterns that can be related to known structural and functional roles.

#### 4.3.3. Δ-Connectome Visualization

For each region–role combination included in the analysis, we identified all valid directed paths in which the region served in the specified role (Source or Target). For each path, we computed the *log*_2_ ratio of efficiency (*E*) between two species:

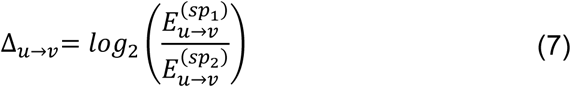

where *sp*_1_and *sp*_2_denote the two species being compared. Positive values indicate higher efficiency in *sp*_1_; negative values indicate higher efficiency in *sp*_2_.

For each partner region connected to the focal region, we averaged Δ values across all relevant paths and retained the top partners ranked by absolute mean delta. A directed subgraph was then constructed with the focal region and its top partners as nodes, edge directions determined by the focal role, node size proportional to ∣Δ∣, node color indicating the sign of Δ, and edge width proportional to ∣Δ∣. Graphs were arranged using a modified circular layout with the focal region centered.

## Acknowledgments

Computational resources were provided by the Pittsburgh Supercomputing Center through NSF ACCESS allocations MED230052 and CIS200026.

## Use of AI Tools

We used ChatGPT (OpenAI) solely for language refinement during manuscript preparation; all scientific content was produced by the authors.

## Author Contributions

*Siva Venkadesh:* Conceptualization, Methodology, Formal analysis, Software, Validation, Visualization, Writing – Original Draft, Supervision

*Yuhe Tian*: Software, Visualization, Writing – Review & Editing

*Wen-Jieh Linn*: Data Curation, Visualization, Writing – Review & Editing

*G. Allan Johnson*: Data Curation, Writing – Review & Editing, Resources, Funding acquisition. *Fang-Cheng Yeh*: Methodology (dMRI tractography), Writing – Review & Editing, Resources, Funding acquisition.

## Code Availability

All analysis and figure-generation code supporting the findings of this study are archived at Zenodo (https://doi.org/10.5281/zenodo.17653309) in the directory *projects/cross_species_connectomics/*.

## Data availability

The mouse viral tracer data are publicly available from the Allen Mouse Brain Connectivity Atlas (https://connectivity.brain-map.org). Derived connectome matrices, region labels, and supplementary tables generated for this study are archived at Zenodo (https://doi.org/10.5281/zenodo.17653309) in the directory *projects/cross_species_connectomics/*.

