## Supplementary Materials for "Directed connectomes across species reveal conserved and divergent pathways of neural signaling"

### Supplementary Material

**Supplementary Table – 1.** Symbol Reference for Equations

| Symbol | Definition | Type / Units |
|---|---|---|
| **Atlas and Masks** | | |
| $A(x)$ | Atlas label at voxel x (0 = background, 1…R = region IDs) | integer |
| $S_k(x)$ | Injection mask for experiment k (binary: 1 = injected voxel, 0 = otherwise) | binary |
| $P_k(x)$ | Projection mask for experiment k (binary: 1 = projected voxel, 0 = otherwise) | binary |
| 1[·] | Indicator function: 1 if condition is true, else 0 | binary |
| **Region-Level Fractions** | | |
| $f_i^{inj,k}$ | Injection fraction for region i in experiment k (Eq. 1) | unitless |
| $f_j^{proj,k}$ | Projection fraction for region j in experiment k (Eq. 2) | unitless |
| **Directed Strengths** | | |
| $w_{i \to j}^{(k)}$ | Experiment-level directed strength from i to j in experiment k (Eq. 3) | unitless |
| $\mathrm{DirMat}_{i,j}$ | Final directed projection strength from i to j, summed over all experiments (Eq. 4) | unitless |
| **Diffusion MRI Measures** | | |
| $L_{ij} = L_{ji}$ | Mean axonal length between regions i and j from tractography | mm |
| $V$ | Total white matter volume for species | $mm^3$ |
| **Effective Costs and Paths** | | |
| $\mathrm{EffCost}_{i \to j}$ | Effective connection cost from i to j (Eq. 5) | unitless |
| $\mathrm{PathCost}_{i \to j}$ | Minimum path cost between i and j (Eq. 6) | unitless |
| $p_{i \to j}$ | Directed path (sequence of edges) from i to j | set of ordered pairs |

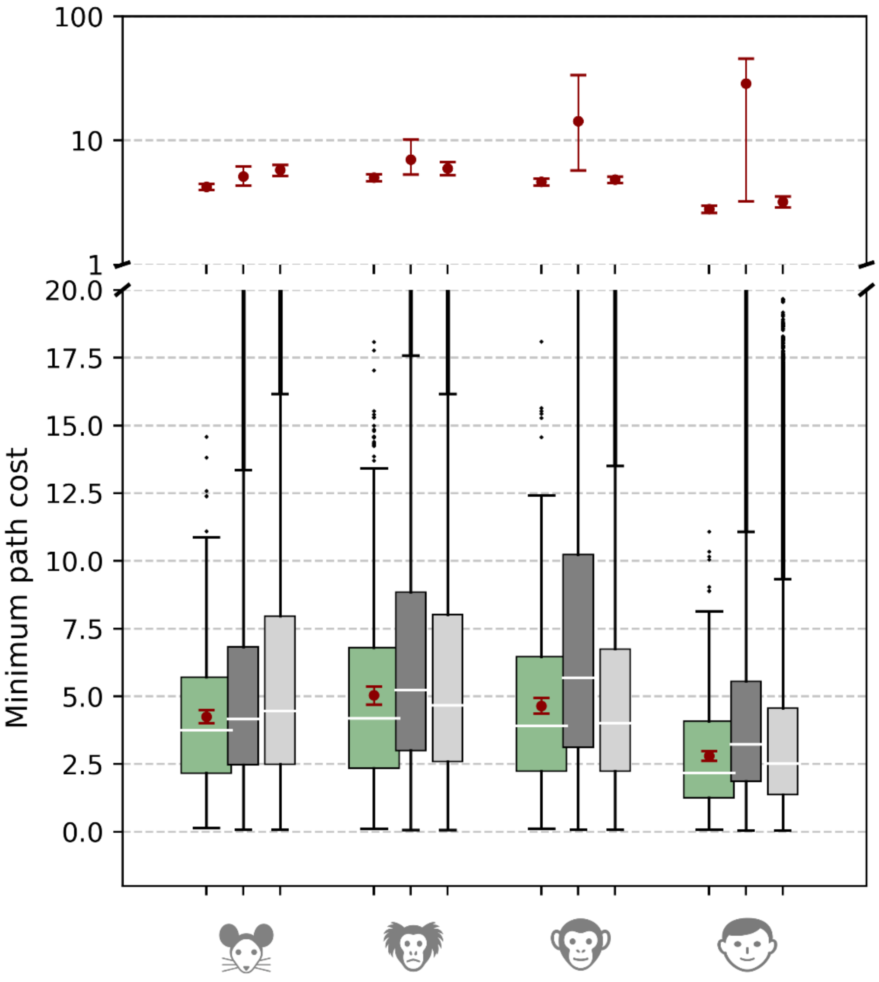


**Supplementary Figure 1 |** Path cost distributions for real connectomes (green) and null networks from 10,000 permutations, shown as the full 462 × 10,000 distributions to illustrate the spread of nulls. Both tracer-directionality shuffles (dark grey) and dMRI-length shuffles (light grey) are shown. Mean path cost (± SD) across null permutations and the corresponding real mean: Mouse: real = 4.25, tracer-shuffle = 5.15 ± 0.48, dMRI-shuffle = 5.79 ± 0.29; Marmoset: real = 5.04, tracer-shuffle = 7.02 ± 1.92, dMRI-shuffle = 5.96 ± 0.38; Rhesus: real = 4.65, tracer-shuffle = 14.27 ± 45.9, dMRI-shuffle = 4.85 ± 0.15; Human: real = 2.80, tracer-shuffle = 28.7 ± 661.6, dMRI-shuffle = 3.21 ± 0.17. Tracer-randomized nulls showed far greater spread than dMRI-length–randomized nulls, underscoring the stabilizing influence of directionality on network organization. Without directional constraints, networks admitted inefficient detours and spurious shortcuts, generating extreme path costs. In contrast, randomizing axonal lengths raised overall costs but retained narrower distributions, consistent with directionality anchoring efficient routes even under altered wiring distances.

**Supplementary Table – 2.** Pairwise Species Comparisons of Regional Efficiency Using Dunn's Test

| Comparison | Z-score | *P* |
|---|---|---|
| **TEM_Inferior (Source)** | | |
| Mouse vs Marmoset | **-3.05** | 0.007 |
| Mouse vs Rhesus | **-3.36** | 0.005 |
| Mouse vs Human | -1.58 | 0.170 |
| Marmoset vs Rhesus | -0.31 | 0.757 |
| Marmoset vs Human | 1.47 | 0.170 |
| Rhesus vs Human | 1.78 | 0.150 |
| **TEM_Superior (Source)** | | |
| Mouse vs Marmoset | 0.03 | 0.972 |
| Mouse vs Rhesus | -1.05 | 0.352 |
| Mouse vs Human | **-3.03** | 0.007 |
| Marmoset vs Rhesus | -1.08 | 0.352 |
| Marmoset vs Human | **-3.07** | 0.007 |
| Rhesus vs Human | -1.98 | 0.095 |
| **INS_Anterior (Source)** | | |
| Mouse vs Marmoset | **3.98** | 0.000 |
| Mouse vs Rhesus | 0.91 | 0.362 |
| Mouse vs Human | -1.8 | 0.086 |
| Marmoset vs Rhesus | **-3.06** | 0.004 |
| Marmoset vs Human | **-5.78** | 0.000 |
| Rhesus vs Human | **-2.71** | 0.010 |
| **INS_Anterior (Target)** | | |
| Mouse vs Marmoset | 2.05 | 0.061 |
| Mouse vs Rhesus | 0.43 | 0.669 |
| Mouse vs Human | **-3.01** | 0.005 |
| Marmoset vs Rhesus | -1.62 | 0.126 |
| Marmoset vs Human | **-5.06** | 0.000 |
| Rhesus vs Human | **-3.43** | 0.002 |
| **OLF_Anterior (Source)** | | |
| Mouse vs Marmoset | -0.92 | 0.356 |
| Mouse vs Rhesus | 1.51 | 0.157 |
| Mouse vs Human | **3.03** | 0.007 |
| Marmoset vs Rhesus | **2.44** | 0.030 |
| Marmoset vs Human | **3.95** | 0.001 |
| Rhesus vs Human | 1.52 | 0.157 |
| **OLF_Piriform (Target)** | | |
| Mouse vs Marmoset | **-3.07** | 0.004 |
| Mouse vs Rhesus | 1.58 | 0.136 |
| Mouse vs Human | 1.88 | 0.090 |
| Marmoset vs Rhesus | **4.66** | 0.000 |
| Marmoset vs Human | **4.96** | 0.000 |
| Rhesus vs Human | 0.3 | 0.766 |

p-values were corrected for FDR.
